# Dissecting the cell of origin of aberrant SALL4 expression in myelodysplastic syndrome

**DOI:** 10.1101/2022.12.05.518121

**Authors:** Hiro Tatetsu, Miho Watanabe, Jun Liu, Kenji Tokunaga, Eisaku Iwanaga, Yoshihiro Komohara, Emily Thrash, Matsuoka Masao, Daniel G. Tenen, Li Chai

## Abstract

Myelodysplastic syndrome (MDS) is a group of heterogeneous diseases characterized by cytologic dysplasia and cytopenias resulting from ineffective hematopoiesis. Oncofetal protein SALL4 is a known oncogene in MDS and its baseline expression level serves as a prognostic biomarker for MDS at the time of diagnosis. In addition, a recent study showed that SALL4 upregulation following hypomethylating agent treatment in MDS patients correlates with poor outcomes. Despite its important mechanistic and diagnostic significance, the cellular identity of bone marrow cells with aberrant SALL4 expression in MDS patients remains unknown.

In this study, we analyzed MDS bone marrow cells on single cell level by mass cytometry (CyTOF) and found that SALL4 was mainly aberrantly expressed in the hematopoietic stem and progenitor cells (HSPC) as well as myeloid lineages. Within the HSPC population from MDS patients, SALL4 and p53 were co-expressed, with the highest co-expressing clones harboring pathogenic TP53 mutations. Overall, our study characterizes for the first time the aberrant SALL4 expression in primary MDS patient samples at a single-cell level. Further studies on the SALL4/p53 network for in-depth mechanistic investigation are needed in the future.

**Key Points:** SALL4 expression in various MDS BM cells confirmed by mass cytometry (CyTOF). SALL4 and p53 double positive cells were predominantly found in the hematopoietic stem and progenitor cell (HSPC) population and associated with pathogenic TP53 mutation status.

## Introduction

Myelodysplastic syndrome (MDS) is a group of heterogeneous pre-leukemic hematological disorders characterized by cytologic dysplasia and refractory cytopenias as a result of ineffective hematopoiesis^1^, as well as increased risk of transformation to acute myelogenous leukemia (AML). As high as 30% of MDS patients progress to AML. MDS is currently diagnosed based on morphological dysplasia upon visual examination of a bone marrow aspirate and biopsy. Additional studies such as karyotyping, flow cytometry and molecular genetics may help to assist in diagnosis. In recent years, there is heightened interest among researchers and clinicians to incorporate molecular biomarkers for risk stratification, which is essential for selecting the appropriate therapeutic interventions. Although our understanding of the pathophysiology of MDS and AML progression improved remarkably in recent years, therapeutic advances have been limited overall. Allogeneic transplantation remains the only cure for MDS at the present time. Therefore, a major effort has been directed towards developing more effective treatments for patients who are ineligible for transplantation^2^.

Although well characterized at the level of cytogenetic changes and gene mutations, the pathogenesis of MDS remains incompletely understood, which poses a major barrier for developing effective therapeutic strategies^2 3^. SALL4 is a known leukemic oncogene and an emerging risk predictor in MDS, particularly in association with hypomethylating agents (HMA) treatment. It is a transcription factor important for development and embryonic stem cell properties. While SALL4 expression is downregulated or absent in most adult tissues, SALL4 is re-expressed in various cancers^4 5^. SALL4 is detectable in the normal CD34+ HSPC population^6^, and the major functional role of SALL4 in normal hematopoiesis is to maintain the stem/progenitor cells in an undifferentiated state with self-renewal capacity. SALL4 is important for histone deacetylase inhibitors (HDACi) mediated hematopoietic stem/progenitor cells (HSPCs) expansion ex vivo^7 8 9^. Studies in murine models, as well as in human samples, have demonstrated that SALL4 is essential for leukemic cell survival and that SALL4 transgenic mice develop an MDS-like phenotype prior to transformation to AML, supporting that SALL4 is an oncogenic driver during the MDS/AML pathogenesis^6^. In a leukemia cell line model, loss of function studies have demonstrated that SALL4 is a key regulator in leukemic cell survival and down-regulation of SALL4 can lead to significant apoptosis of the cells^10^. SALL4 has a negative impact on DNA damage repair, and a model of dual functional properties of SALL4 in leukemogenesis by inhibiting DNA damage repair and promoting cell survival has been proposed^11^.

In MDS patients, expression of SALL4 was increased and correlated with disease progression^12^. Recently, a multi-center cohort study reported that demethylation and upregulation of SALL4 could occur in MDS patients treated with hypomethylating agents, a class of drugs to which half of MDS patients show primary resistance and nearly all develop secondary resistance. Importantly, SALL4 upregulation was found to be associated with a worse survival outcome and this study addressed a major question in terms of mechanisms of HMA failure^13^. Alarmingly, at the present time, there are no approved interventions for MDS patients with progressive or refractory disease, particularly after HMA-based therapy. Although an FDA approved SALL4-targeting drug does not currently exist, several promising drug leads are being developed^14^, and SALL4-centered therapy in combination with HMA for MDS/AML patients is under active investigation. SALL4 may have important relationships with other key players in MDS pathophysiology. For example, we previously reported that higher SALL4 mRNA expression in MDS patients was associated with the complex karyotype and its expression was positively correlated with p53 expression in MDS patients by immunohistochemistry^6 11^. TP53 mutations are identified in over 70% of MDS patients with complex karyotype^15^, and confer poorer prognosis^16^.

Although a series of recent studies demonstrated SALL4’s crucial roles in the pathogenesis and treatment outcomes in MDS, several key questions remain to be addressed. First of all, the relationship between SALL4 expression with somatic gene mutations (e.g. TP53), which are important for risk stratification and identification of disease drivers, hasn’t been well explored in MDS patients. Furthermore, the identity of SALL4-expressing cells in the bone marrow of MDS patients is unknown. Dissecting the cell of origin for SALL4’s action has important implications for understanding MDS pathogenesis, as well as SALL4 therapeutic targeting in myeloid neoplasms.

Mass cytometry, also called cytometry by time of flight (CyTOF^®^), combines key aspects of flow cytometry with distinct advantages of mass spectrometry. This technique enables simultaneous detection of over 40 parameters per cell for up to millions of cells. Mass cytometry has significantly increased our ability to profile entire populations of cells at the individual cellular level. Using single cell mass cytometry and paired bone marrow Whole Exome Sequencing (WES), we for the first time comprehensively evaluated the expression of SALL4 in relation to other oncoproteins and gene mutations in bone marrow cells from patients with MDS. We identified the cell of origin of aberrant SALL4 expression in the HSPCs and myeloid lineages, as well as a strong correlation between aberrant SALL4 and TP53 expression in the HSPCs. While this study serves as a starting point to assess the pathogenic roles of SALL4 in MDS on a single cellular level, we also demonstrated the ability of the CyTOF technology to capture intratumor heterogeneity of SALL4 expression, which serves as a proof-of-principle study to further the diagnostic development for oncoproteins such as SALL4 that is aberrantly expressed in only select lineages but with important prognostic/therapeutic implications.

## Methods

### Study approval

This research was approved by the Ethics Committee of Kumamoto University (approval genome No. 297). DNA from bone marrow mononuclear cells from 10 MDS was extracted using the according to the manufacturer’s recommendations (**Suppl Table 1)**. We performed CyTOF^®^ (Fluidigm^®^) and WES (Novogene Bioinformatics Institute, Beijing, China) for MDS patients and CyTOF for 5 lymphoma patients, on which the invasion of bone marrow was ruled out, as the controls.

### Metal-tagged monoclonal antibodies

A panel of 27 metal-tagged monoclonal antibodies was used for analysis of the patient’s bone marrow mononuclear cells. A detailed listing of antibodies and corresponding metal tags is provided in **Suppl Table 2**. Antibodies were purchased either already metal-tagged (Fluidigm®) or in purified form. They were labeled using the Maxpar® X8 Antibody Labeling Kit (Fluidigm) according to the manufacture’s recommended protocol, titrated, and were diluted to 0.5 mg/ml in Antibody Stabilizer (CANDOR Bioscience) for long-term storage at 4°C.

### Antibody staining for mass cytometry

Bone marrow mononuclear cells were thawed at 37°C, resuspended in 10% FBS in RPMI1640 and washed twice with PBS (without calcium or magnesium). Bone marrow blood cells were then transferred to 1.5ml tube and were stained 5 minutes in PBS supplemented with 0.2μM Cisplatin Cell-ID™ (Fluidigm, San Francisco, CA). Cells were suspended and washed with Maxpar® cell staining buffer (Fluidigm). Then cells were resuspended in Maxpar® cell staining buffer (Fluidigm) and blocked with Human TruStain FcX™(Biolegend) for 10 minutes at room temperature. Cells were incubated with an antibody targeting cell surface markers for 30 minutes at room temperature and then washed twice with cell staining buffer. After washing, cells were fixed and permeabilized for 30 minutes on ice using eBioscience FoxP3 fix/perm (Thermo Fisher Scientific, Waltham, MA, USA). Fixed/permeabilized cells were washed twice with 1×working solution of permeabilization buffer(eBiosciences) and incubated with all antibodies targeting intracellular antigens for 30 minutes at room temperature. After staining with intracellular antibodies, cells were washed twice with 1× working solution of permeabilization buffer and incubated with Cell-ID Ir DNA intercalator (Fluidigm) over night at 4°C. On the next day, prior to mass cytometry analysis, cells were washed twice with Maxpar® cell staining buffer (Fluidigm) and resuspended in water containing EQ™ Four Element Calibration Beads (Fluidigm). Samples were acquired on a Helios™ CyTOF System (Fluidigm).

### Mass cytometry analysis

Cells were analyzed on a mass cytometry (Helios, CyTOF System) (Fluidigm) at an event rate of approximately 300 cells/second. The resulting data were analyzed with software available through Cytobank (www.cytobank.org). For mass cytometry data, all parameters except time and event length were displayed with an arcsinh transformation and a scale augment of 5 ranging from −5 to 12000. Event length and time were displayed on linear scales. 191Ir and 193Ir DNA intercalator and 140Ce beads were used to discern intact singlets from debris and cell aggregates. Then live single cells were selected by applying a gate on 193Ir DNA vs. 198Cisplatin. All gating and extraction of median expression levels were performed using Cytobank. To interpret high-dimensional single-cell data that were produced by mass cytometry, we used a visualization tool based on the viSNE and FlowSOM algorithm, which allows visualization of high-dimensional cytometry data on a 2-dimensional map at single-cell resolution and preserves the nonlinearity^17 18^. FlowSOM is an algorithm that allows us to visualize both manual gatings and automated clusterings to divide cell types into clusters and meta-clusters, based upon the similarity of cell-specific marker expression profiles using pie charts ^18^. Immunophenotypic subsets on the basis of standard surface markers were analyzed based on the European LeukemiaNet recommendation and the related publications ^19 20 21^ **(Suppl Table 2)**. All gating and extraction of median expression levels were performed using Cytobank. CyTOF data analysis was performed using viSNE and FlowSOM, which is an advanced clustering analysis software in Cytobank™ (Beckman Coulter Life Sciences, Indianapolis, IN).

Cell subsets defined as HSPC (haematopoietic progenitor stem cells): Lin-CD34+, HSC (haematopoietic stem cells): Lin−CD38−CD34+, MPP (multipotent progenitors): Lin−CD34+CD38+CD90−CD45RA-CD49f−, CMP (common myeloid progenitors): Lin−CD38+CD34+ CD90−CD123+CD45RA−: GMP (granulocyte macrophage progenitors): Lin−CD38+ CD90−CD34+CD123+CD45RA+ and MEP (megakaryocyte-erythroid progenitors): Lin−CD38+ CD90−CD34+CD123+CD45RA+cells. Lineage negative cells were defined as following CD11b, CD38, CD3, CD7, CD19, CD71, CD235, and CD33. Monocyte Lineage cells were defined as following criteria: not CD34+CD38 low, CD33 positive, HLA-DR positive, not CD123 bright and CD33 low, CD3 negative, not CD19+, not CD45 high and CD7 high, not brightly CD38 positive, not CD71 high or CD235 high. Granulocyte Lineage cells were defined as following criteria: not CD71 high or CD235 high, CD3 negative, not CD19+, not brightly CD38 positive. Erythroid Lineage cells were defined as following criteria: not CD34+CD38 low, CD3 negative, not CD19+. Within this population, CD71 bright CD235 low cells were defined as pro-erythroblasts, CD71 bright CD235 positive cells were defined as early erythroblasts, and late erythroblasts were defined as CD235 positive CD71 mid and CD45 negative. B cell Lineage cells were defined as following criteria: CD33 low, CD3 negative, not CD71high or CD235 high, CD11b low. CD3 positive was defined as T cells. NK cells were defined as following: positive for CD7 and CD45, expression of CD7 in the absence of CD3, and lack of CD34 expression.

### Data mining

Publicly available database GSE19429 consists of gene expression profiles from MDS patients of various subtypes. Gene expression profiles were generated using Affymetrix Human Genome U133 Plus 2.0 Array. The serial matrix files were downloaded from Gene Expression Omnibus.

### Whole Exome Sequencing

The bone marrow mononuclear cells were collected using Ficoll-Paque™ PREMIUM (Cytiva, Tokyo, Japan). Novogene (Beijing, China) performed the whole-exome sequencing (WES) including exome capture, high throughput sequencing, and common filtering by purified genomic DNA. WES data were analyzed for somatic single nucleotide polymorphisms (SNP), insertions and deletions (INDEL). Analyses were conducted of the relevant mutations of 62 genes, listed in **Suppl Table 3**.

### Statistical analysis

All statistical analyses were performed with Excel and EZR (Saitama Medical Center, Jichi Medical University, Saitama, Japan), which is a graphical user interface for R (The R Foundation for Statistical Computing, Vienna, Austria). More precisely, it is a modified version of R commander designed to add statistical functions frequently used in biostatistics^22^. Statistical significances of two group comparisons were determined using Mann–Whitney U test. Multiple comparisons were performed by one-way ANOVA with Tukey correction. The minimum significance level was set at *P* < 0.05. Asterisks indicate the statistical significance as follows: \**P* ≤ .05; \*\**P* ≤ .01.

## Results

### The mRNA expression of SALL4 was increased in CD34+ cells in MDS patients

Given the known functions of SALL4 in HSPCs, we first examined the expression of SALL4 mRNA in MDS CD34+ cells by analyzing MDS expression profiles from the public database GSE19429, which was obtained from 183 MDS patients and 17 healthy controls^23^, we noticed that SALL4 expression in MDS CD34 cells was higher than that of controls, and the highest in refractory anemia with excess blasts type 2 (RAEB2) (*P* ≤ .05) **(Figure 1A & B)**.

**Figure 1.**
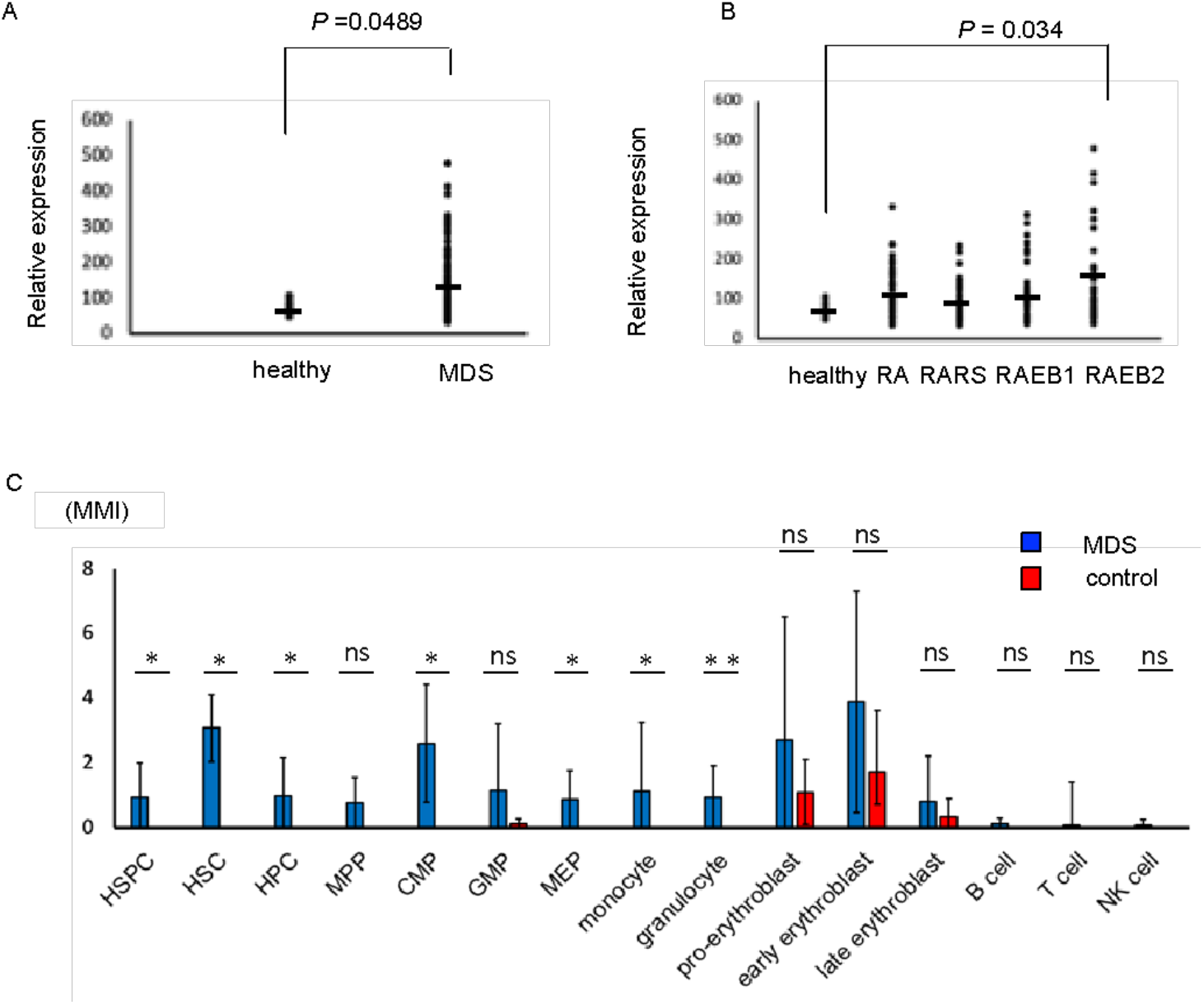
SALL4 RNA and protein were expressed in MDS bone marrow cells. (A) SALL4 expression in MDS CD34 positive cells from the public database GSE19429. (B) SALL4 expression in CD34 positive cells in MDS subtype from the public database GSE19429. abbreviations: RA, refractory anemia; RARS, refractory anemia with ring sideroblasts; RAEB1, refractory anemia with excess blasts1; RAEB2, refractory anemia with ring sideroblasts2. (C) MMI of SALL4 in bone marrow cells on CyTOF (MDS patients n=10, control BM n=5, Error bars indicate standard deviation). Data analysis used Mann–Whitney U test. \**P* ≤ .05; \*\**P* ≤ .01. Abbreviations: ns, not significant; HSPC, hematopoietic stem and progenitor cell; HSC, Hematopoietic stem cell; HPC, hematopoietic progenitor cell; MPP, multipotent progenitors, CMP; common myeloid progenitor; GMP granulocyte-monocyte progenitor; MEP, megakaryocyte-erythrocyte progenitor; NK cell, natural killer cell; MMI, median metal intensity.

### Aberrant SALL4 protein expression is detected in HSPC and myeloid lineage in MDS BM samples

To examine the protein expression pattern of SALL4 in MDS patients’ BM, we performed CyTOF experiments in 10 MDS patients as compared with 5 control BM samples. The staining panels included 22 surface markers of cell surface proteins and 5 markers of intracellular signaling **(Suppl Table 2)**. We first validated the CyTOF method detection of SALL4 using NB4, a SALL4 positive human promyelocytic leukemia cell line, and MT4, a SALL4 negative human adult T-cell leukemia cell line **(Suppl Figure 1)**^10^. For population analysis, we used the manual gating approach based on the existing publications^21^. We observed that MDS patients had aberrant SALL4 protein expression in hematopoietic stem cells (*P* ≤ .05), hematopoietic progenitor cells (*P* ≤ .05) and myeloid lineage cells (*P* ≤ .05) when compared to controls (**Figure 1C & Suppl Figure 2)**. We also noticed that within the same MDS patient BM sample, SALL4 protein was expressed in myeloid and erythroid lineages but absent in lymphoid lineage. In addition, SALL4 protein expression could be detected in erythroid lineages in control BM cells as well.

**Figure 2.**
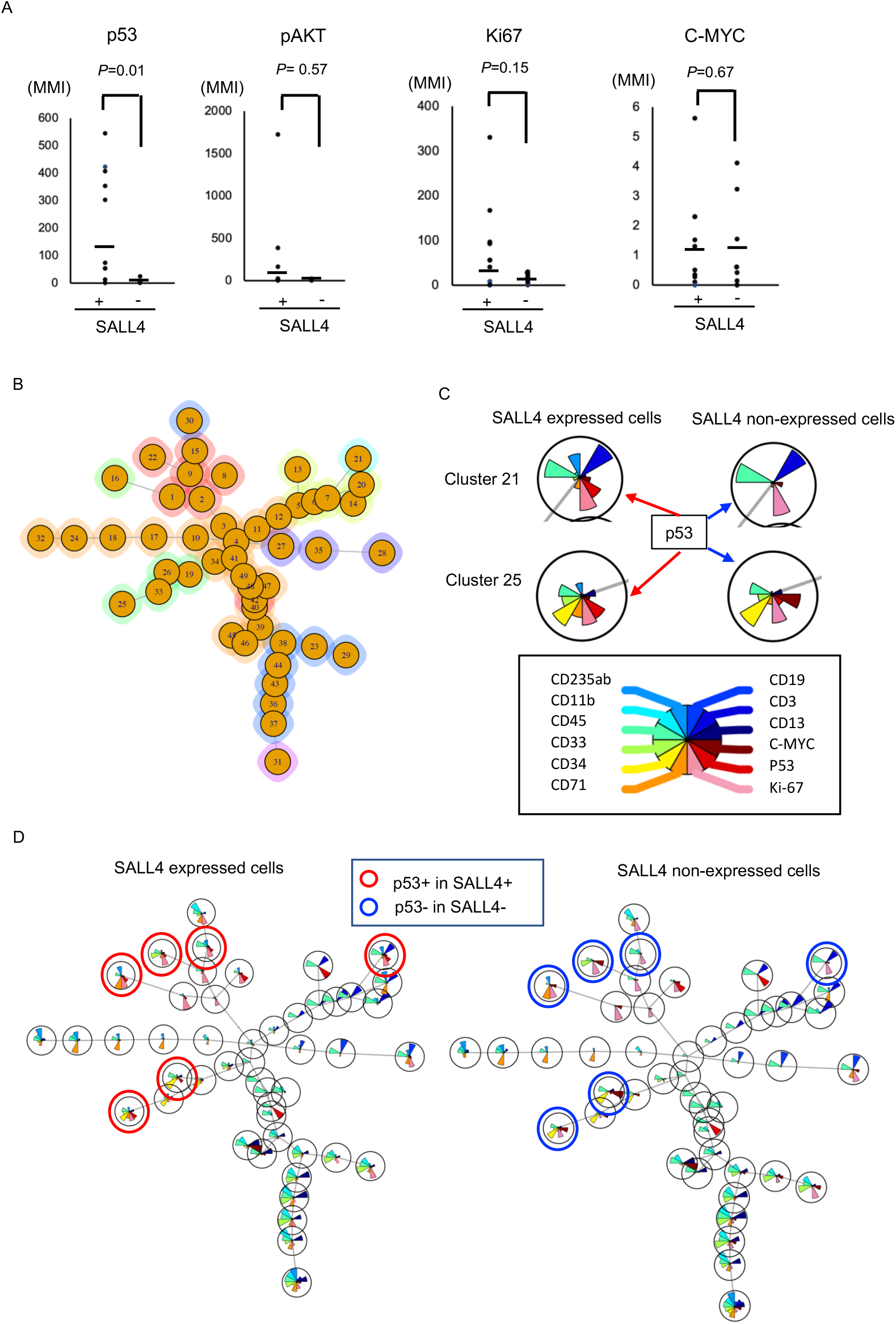
ViSNE analysis on SALL4 expressing cells in MDS samples. (A) P53, ki67, c-myc or pAKT expression of SALL4 expressing cells and SALL4 non-expressing cells. (B) Cells were clustered into 49 groups on FlowSOM analysis. (C) The example of FlowSOM via viSNE analysis based on the distribution of SALL4, p53, Ki67 and major lineage markers (CD235ab, CD11b, CD45, CD33, CD34, CD71, CD19, CD3, CD11b) on SALL4 expressing cells (left side) and SALL4 non-expressing cells (right side) on cluster 21 and 25. The brown fan shape indicates p53. (D) FlowSOM via viSNE analysis on SALL4 expressing cells (left panel) and SALL4 non-expressing cells (right panel). The pie-chart visualizes the relative expression. The red circle indicated that p53 expressing cells in SALL4 expressing cells. The blue circle indicated that p53 non-expressing cells in SALL4 non-expressing cells, which were a pair of p53 expressing cells in SALL4 expressing cells.

### SALL4 expression was correlated with p53 expression at the single cell level in the HSPCs

Next, we took advantage of CyTOF that offers protein expression at single cell level, and investigated the relationship between SALL4 expression with p53, ki67, c-myc or phosphorylated AKT (pAKT), which are the proteins among important pathways in leukemia or MDS pathogenesis^15,24-26^. The bone marrow cells in the individual patients were divided into SALL4 expressing cells and SALL4 non-expressing cells, and we analyzed the expression levels of these factors for each group. SALL4 expressing cells also expressed p53 (*P*=0.01) in HSPC **(Figure 2A)**. SALL4 expressing cells also showed a trend of a higher level of pAKT and ki67, although the differences are not statistically significant. TP53 mutations are adverse prognostic factors in a wide variety of clinical settings^27,28^. Although Immunohistochemistry can’t normally detect the wild-type (WT) p53 protein due to its short half-life, mutated proteins are usually detectable due to prolonged half-life^29^.

To understand the expression pattern of SALL4 and p53 and dissect the relationship between SALL4 and p53 in MDS patients, we performed FlowSOM via viSNE analysis on MDS bone marrow cells, which were divided into two groups: those without SALL4 expression and those with SALL4 expression. FlowSOM analysis distinguished 49 clusters, based on the expression of p53, Ki67 and the major lineage markers (CD235ab, CD11b, CD45, CD33, CD34, CD71, CD19, CD3, CD11b) **(Figure 2B&C)**. SALL4 expressing clusters also tended to express p53, conversely, SALL4 non-expressing cells were not **(Figure 2C&D)**.

Next, we performed the minimal spanning trees (MSTs) analysis by using FlowSOM in MDS patients **(Figure 3A)**. FlowSOM distinguished 10 meta-clusters by using CD235, CD11b, CD45, CD33, CD34, CD71, CD19, CD10, CD3, CD13, p53 and SALL4. Based on these clusters, we analyzed SALL4 and/or p53 expressing or non-expressing cells **(Figure 3B, Suppl Figure 3A)**. Most of the cells did not express SALL4 and p53, which were mainly included in meta-cluster 3 **(Suppl Figure 3B)**. SALL4+p53-cells were included in meta-cluster 2**(Suppl Figure 3C&4)**. While SALL4-p53+ cells were included in C3, C9 and C10, SALL4+p53+ cells were exclusively included in meta-cluster 9. **(Figure 3C, Suppl Figure 3D&4)**.

**Figure 3.**
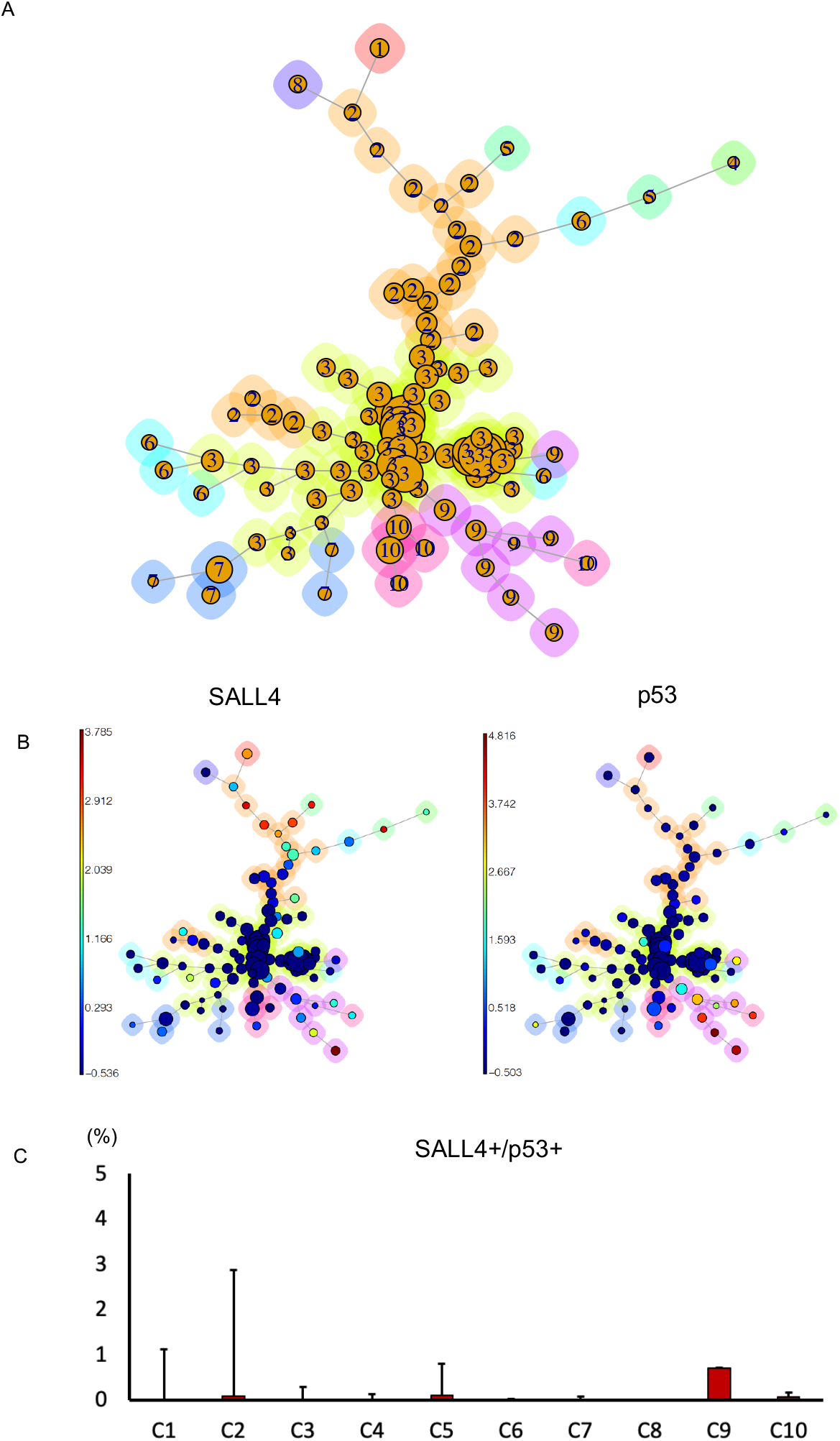
Examination of SALL4 and p53 expression in MDS bone marrow cells. (A) Cells were clustered into 12 nodes and depicted as a minimal spanning tree, to the ten identified clusters on FlowSOM analysis. (B) Aggregated events of SALL4 (left) and p53 (right) on FlowSOM analysis. (C) The percentage of SALL4+p53+ cells. Error bars indicate standard deviation.

Interestingly, both patients NO.1 and NO.2, who have the highest fractions of SALL4/TP53 double positive cells (meta-cluster 9: 56.3% and 40%, respectively), harbored pathogenic loss-of-function TP53 mutations in the DNA binding domain **(Table 1)**. The c.476C>T mutation of patient NO.1, is a missense variant that occurs at a mutational hotspot within TP53 (cBioPortal). A combination of *in vitro* and in silico data suggests that this variant is deleterious to TP53 function (IARC TP53 database). This mutation is predicted to be pathogenic (COSMIC), and when occurring in the germline, has been recently found to cause Li-Fraumeni syndrome ^31^. Patient NO.2 harbors a splice site variant, c.782+1G>T, which alters the canonical splice donor site for intron 7. It has been reported as a somatic variant in thymic cancer ^32^ as well as a pathogenic variant in Li-Fraumeni syndrome (ClinVar). For patient NO.7, the p.V173* mutation is a frameshift variant at a minor mutational hotspot of the DNA binding domain, with many pathogenic missense variants reported to date (cBioPortal, COSMIC). Although this particular variant has not been reported in major mutational databases, it likely results in nonsense mediated decay and reduced/absent TP53 protein expression. Thus, it’s not surprising that a significantly smaller fraction of SALL4/TP53 positive cells is detected in patient NO.7 despite the presence of a deleterious TP53 mutation, as compared to patients NO.1 and NO.2.

**Table 1.** SALL4 & p53 double positive cells and karyotypes & mutations in MDS.

| Sample number | Cytogenetics | Mutations | % of SALL4 & p53 DP cells (C9) | TP53 mutation s, if present | Variant description |
| --- | --- | --- | --- | --- | --- |
| 1 | 44,X,-Y,-5,-7,8,-9,add(17)(p11.2),-18,-20+mar1,mar2,mar3[12]/46,XY[3] | GNAS, GATA2, KMT2C, TP53 | 56.3 | c.476C>T | Missense variant in DNA binding region |
| 2 | 46,XY,-6,-17,-19,-20,+mar1,+mar2[15]/46,XY[5] | KMT2C, TP53 | 40 | c.782+1G>C | Intronic splice site variant, DNA binding domain |
| 3 | 46,XX,+1,der(1;7)(q10;p10){11}/45, idem, -18[4]/46,XX[5] | KMT2C | 22.5 | - | - |
| 4 | 46,XX,t(3;21)(q26.2;q22.1)[20] | CALR, GATA2, KIT | 14.3 | - | - |
| 5 | 46,XY,der(1;7)(q10;p10)[1]/46,XY[19] | ETNK1, FLT3, KMT2C, TET2, ZRSR2, | 13.02 | - | - |
| 6 | 46,XY,+1,der(1;7)(q10;p10)[6]/idem,del(20)(q1?)[11]/46,XY[3] | ETNK1, EZH2, GATA2, PTPN11, SETBP1, TERT | 9.54 | - | - |
| 7 | 42,XY,-4,-5,-7,add(7),9,add(12)(p11.2),add(12)(q24.1),add(13)(p11.2),-17,-18,-20,+mar1,+mar2,+mar3[15]/46,XY[3] | MYBL2, TP53 | 6.74 | p.V173* | Frameshift variant affecting DNA binding domain |
| 8 | 46,XX, del(20)(q1?)[5]/46,XX[15] | DNMT3A, GATA2, GNAS, SF3B1 | 5.33 | - | - |
| 9 | 46,XY | DNMT3A, ETNK1, SF3B1, TET2 | 3.68 | - | - |
| 10 | 46,XY | PRPF8, RUNX1, SF3B1, TET2 | 2.43 | - | - |

Abbreviations: DP, double positive

## Discussion

Using the state-of-art single cell mass cytometry and paired bone marrow Whole Exome Sequencing (WES), this study for the first time identified the aberrant expression of SALL4 in hematopoietic stem cells, hematopoietic progenitor cells and myeloid lineage cells of MDS BM cells. In addition, we observed a significant SALL4+p53+ cluster in the MDS bone marrow, which is especially prominent in patients harboring pathogenic TP53 mutations.

Despite being an important leukemia oncogene, SALL4 expression in myeloid malignancy has only been analyzed at the level of whole bone marrow samples until now. SALL4 is a known oncogene in myeloid malignancies. In a mouse model with aberrant SALL4 expression, mice showed myelodysplastic syndrome–like features and subsequently leukemic transformation through activation of the Wnt–beta-catenin pathway^30^. In addition, we have previously shown that SALL4 has a negative impact on DNA damage repair, and supports the model of dual functional properties of SALL4 in leukemogenesis through inhibiting DNA damage repair and promoting cell survival^11^. Understanding the cell of origin of SALL4 is not only important for elucidating MDS pathogenesis, but is also important for designing appropriate therapeutic interventions targeting the SALL4-positive malignant clones. Recent advances in the understanding of AML pathogenesis and refined analysis of AML bone marrow cell-of-origin studies have led to the design of therapeutic targeting of leukemia stem cells as the more promising approach toward a cure^31^. Meanwhile, exciting new development of SALL4-targeted therapeutic leads, such as immunomodulary drugs (IMiDs), proteolysis-targeting chimaeras (PROTACs), and molecular glues, are underway^14^.

As a fetal-oncogene, SALL4 expression is significantly regulated by epigenetic modifications, such as DNA methylation. A critical region of CpG island at SALL4 locus that can be affected by the de-methylation therapy (hypomethylating agents, or HMA) is important for SALL4 expression, and the increase of SALL4 expression is strongly predictive of poor patient outcomes following HMA treatment^13^. Our previous and current study have shown that MDS patients with aberrant SALL4 expression tended to have abnormal karyotypes, which is known to confer poorer prognosis in patients with MDS **(Table 1)**^11^. It is plausible that there is an association between global disruption of the methylation landscape with chromosomal instability. For example, a recent study analyzed genome-wide DNA methylation in 649 AML patients and identified a configuration of distinct 13 subtypes (termed “epitypes”). Epitypes E12 and E13 lacked a consistent mutation pattern but retained mutations associated with genomic instability, such as *TP53* mutations and complex karyotype ^32^. It will be interesting to examine the overall methylation landscape, and especially methylation status on the critical region of CpG island at the SALL4 locus in MDS patients with complex karyotypes. However, due to the limited materials from our MDS patient samples, future studies are needed to further explore this hypothesis.

In this study, we observed significant single-cell level co-localization of SALL4 and p53 in a cohort of MDS patients who mostly harbor abnormal karyotypes. TP53 mutations have highly adverse prognostic implications in a wide variety of clinical settings and is considered an independent risk predictor in MDS^27,28^. Clonal cytogenetic abnormalities are detected in 30-50% of de novo MDS cases, and more than 80% of therapy-related cases^33^. Mutations in the TP53 gene are identified in over 70% of MDS patients with complex karyotype, defined as three or more somatic chromosomal abnormalities present in a single clone^15^. Patients with at least two hits of TP53 showed higher genome instability, and unsurprisingly chromosomal abnormalities^34^. Our observation is consistent with a previous report on the level of whole bone marrow samples, showing that higher SALL4 expression was associated with the complex karyotype and higher TP53 expression in MDS patients by immunohistochemistry staining^11^. Nonetheless, the ability to confirm colocalization between these two key MDS pathogenic/prognostic factors on a cellular level raises interesting questions of synergy between TP53 and SALL4 and clonal advantage of the double-positive cells.

The mechanistic interplay between SALL4 and TP53 await further investigation. Recently, the Ebert group used a CRISPR-Cas9 system to generate isogenic human leukemia cell lines with the most common TP53 missense mutations^35^. They found that the hotspot pathogenic mutations in the DNA binding domain led to the loss of function and exert a dominant negative effect, which appears to drive the selective advantage of TP53 mutant clones in myeloid malignancies. We have reported that SALL4 contributes to leukemogenic processes by allowing accumulation of mutations in cells, which is through inhibiting DNA damage repair and promoting cell survival with mutations^11^. The concomitant high SALL4 expression in these TP53 mutant cells may represent a new SALL4/mutant TP53 network and may confer additional advantages and oncogenic potentials. Further mechanistic and functional studies are needed to elucidate whether a synergy exists between SALL4 and TP53 mutants in promoting MDS tumorigenesis. The relationship between SALL4 and wild-type (WT) P53 in cancer is less clear. Recently, Wang et al demonstrated that murine Sall4 exerts its anti-apoptotic function in a p53-dependent manner in the ES cells^36^. Future studies should investigate whether this interaction stays intact or is disrupted in MDS patients.

In addition to offering future directions of mechanistic investigations and therapeutic targeting, this study also serves as a proof-of-principle to further the diagnostic CyTOF development for oncoproteins such as SALL4 that are aberrantly expressed in only select lineages but with important prognostic/therapeutic implications. Risk stratification is critical for guiding treatment for MDS. The conventional prognosis and risk stratification incorporates peripheral cytopenias, percentage of blasts in the bone marrow and cytogenetic characteristics (e.g. IPSS-R)^37^. Recently, increasing effort and attention have been directed towards incorporating additional molecular signatures for risk stratification. A recent study integrated IPSS-R and mutation scores to formulate a machine-learning guided novel risk stratification system, which was shown to be more effective on prognosis and treatment guidance for MDS patients^38^. Some oncogenes, such as SALL4 and many cancer-germline genes, are infrequently mutated but are often dysregulated at the epigenetic level leading to altered levels of protein expression^39^. Mass cytometry combines key aspects of flow cytometry with distinct advantages of mass spectrometry, allowing the profiling of up to 40 protein markers, at the individual cellular level. Although only a few hematopoietic malignancies such as AML and multiple myeloma have been more extensively investigated *via* CyTOF, a few other studies using CyTOF for MDS bone marrow samples have shown promises of the technique in facilitating the identification of disease biomarkers that have the potential to aid in the diagnosis and prognostication of MDS^40^. While our study is underpowered to do so, future studies with larger cohorts can further explore the clinical utility of combining mutational analysis with expression profiling for MDS diagnosis, prognostication, and predicting response to therapy.

In summary, our study has demonstrated for the first time the cell-of-origin of the aberrant expression of fetal-oncoprotein SALL4 at the HSPC level and the myeloid lineage of MDS BM cells through CyTOF technology. In addition, we observed a SALL4+p53+ cluster in MDS patients, which is especially prominent in patients harboring pathogenic TP53 mutations. With the established CyTOF method to detect SALL4, we plan to delineate the mechanism(s) and functional relationship between SALL4 and p53, as well as to evaluate the role of a SALL4-targeted therapy and diagnostics for MDS patients in the future.

## Acknowledgment

This work was supported in JSPS KAKENHI Grant Number 17K09930 and 21K08420 (To H.T). This work was also supported by Singapore Ministry of Health’s National Medical Research Council (Singapore Translational Research (STaR) Investigator Award, D.G.T.; NMRC/OFIRG/0064/2017.); Singapore Ministry of Education under its Research Centres of Excellence initiative; NIH/NCI Grant R35CA197697 and NIH/NHLBI P01HL131477 (D.G.T); as well as Xiu research fund (L.C.). We acknowledge Kiyota Akifumi for assistance of CyTOF and Miho Matsumoto and Hiroto Takeya for technical assistance.

## Author contributions

H.T. designed and performed research, interpreted the data and wrote the paper; M.W performed CyTOF and interpreted data; T.K, H.T and Y.K made a provision of study materials or patients; E.I performed WES and interpreted data; E.T, J.L M.M and D.T. was responsible for critical reading of the manuscript and important intellectual content; and L.C. was responsible for the study concept, design and execution of the research, interpretation of data, and writing and revising the draft paper.

