## Supplemental figures for "Dissecting the cell of origin of aberrant SALL4 expression in myelodysplastic syndrome"

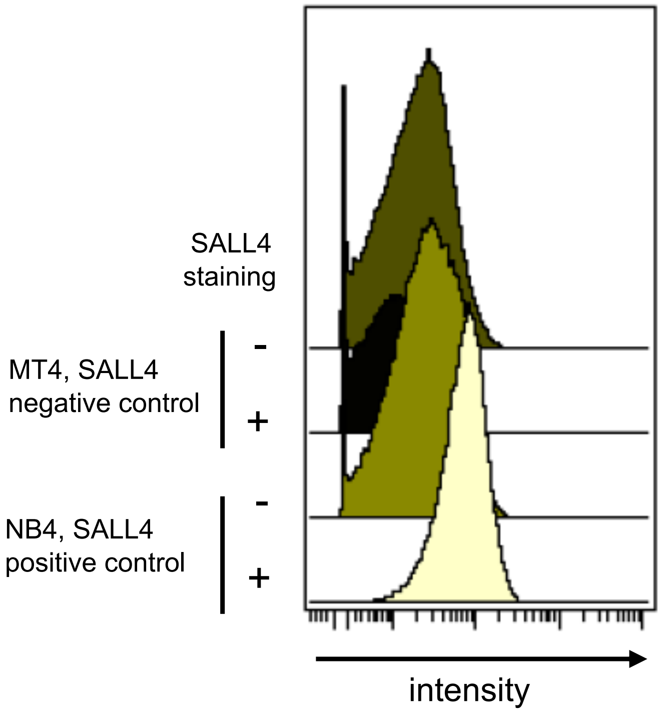


**Supplementary Figure 1. Detection of SALL4 protein in leukemia cell lines via CyTOF**

SALL4 was expressed in NB4, a human promyelocytic leukemia cell line and was not expressed in MT4, a human adult T-cell leukemia cell line.

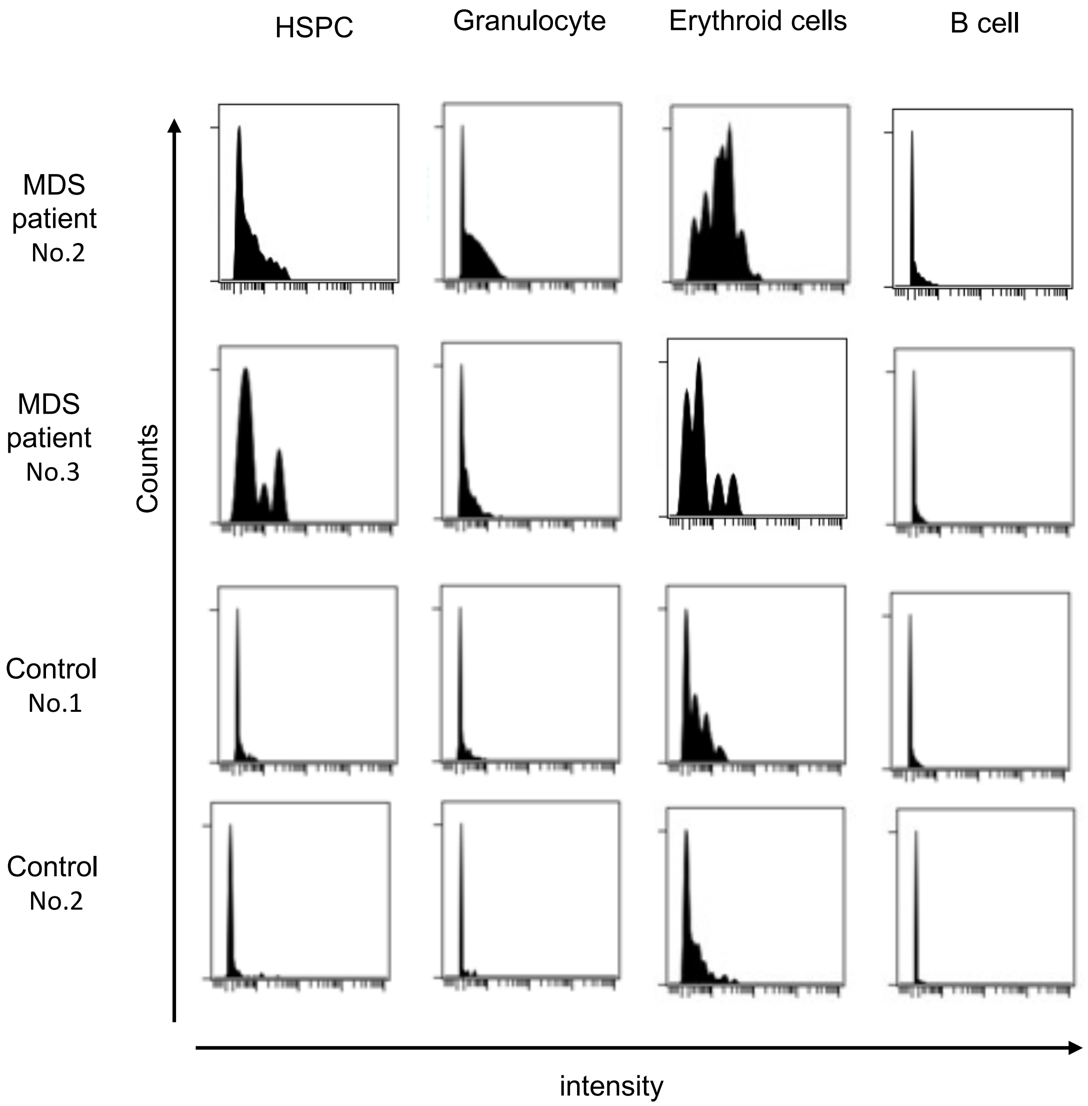


**Supplementary Figure 2. Representative CyTOF data of SALL4 expression in MDS patients and controls**

The upper two panels show SALL4 expression in HSC, granulocyte, B cell and erythroid cells in two representative MDS patients (No.2 and No.3). The lower two panels show those in two representative control patients. Abbreviations: HSPC, hematopoietic stem/progenitor cells.

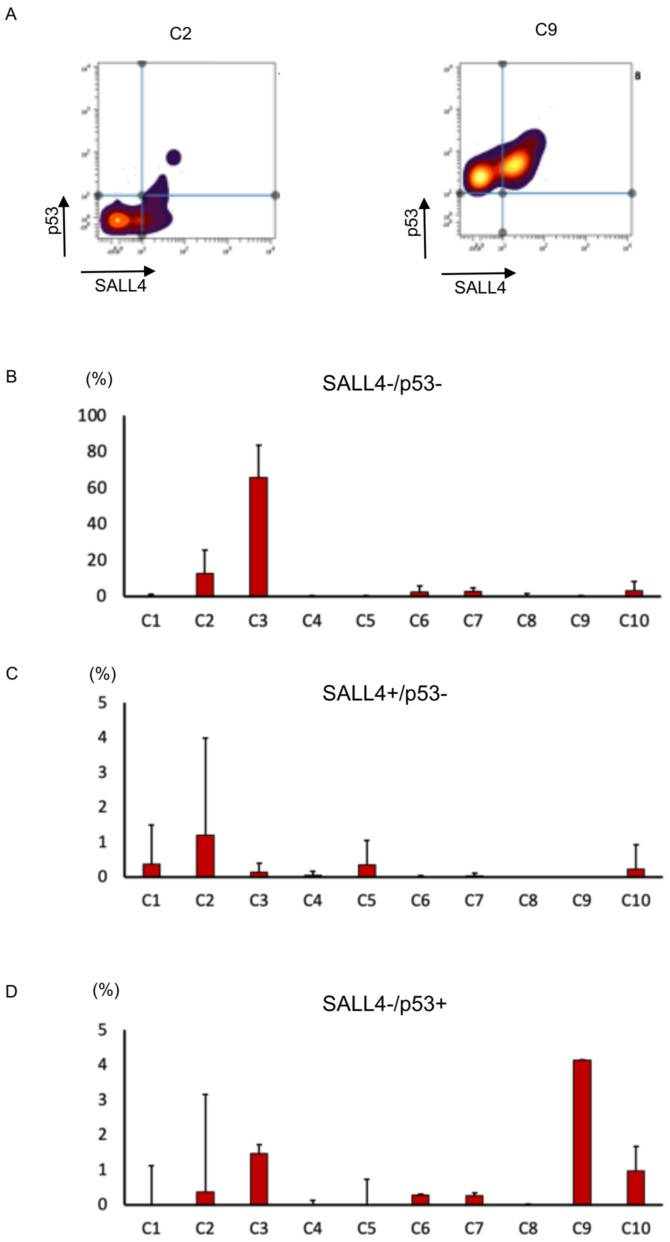


**Supplementary Figure 3. Further analysis on SALL4 and/or p53 expressing cells on MDS samples**

(A) The panel shows a representative pattern of Cluster2 (C2) and Cluster9 (C9) on CyTOF. (B) The percentage of SALL4-p53- cells. (C) The percentage of SALL4+p53- cells (D) The percentage of SALL4-p53+ cells.

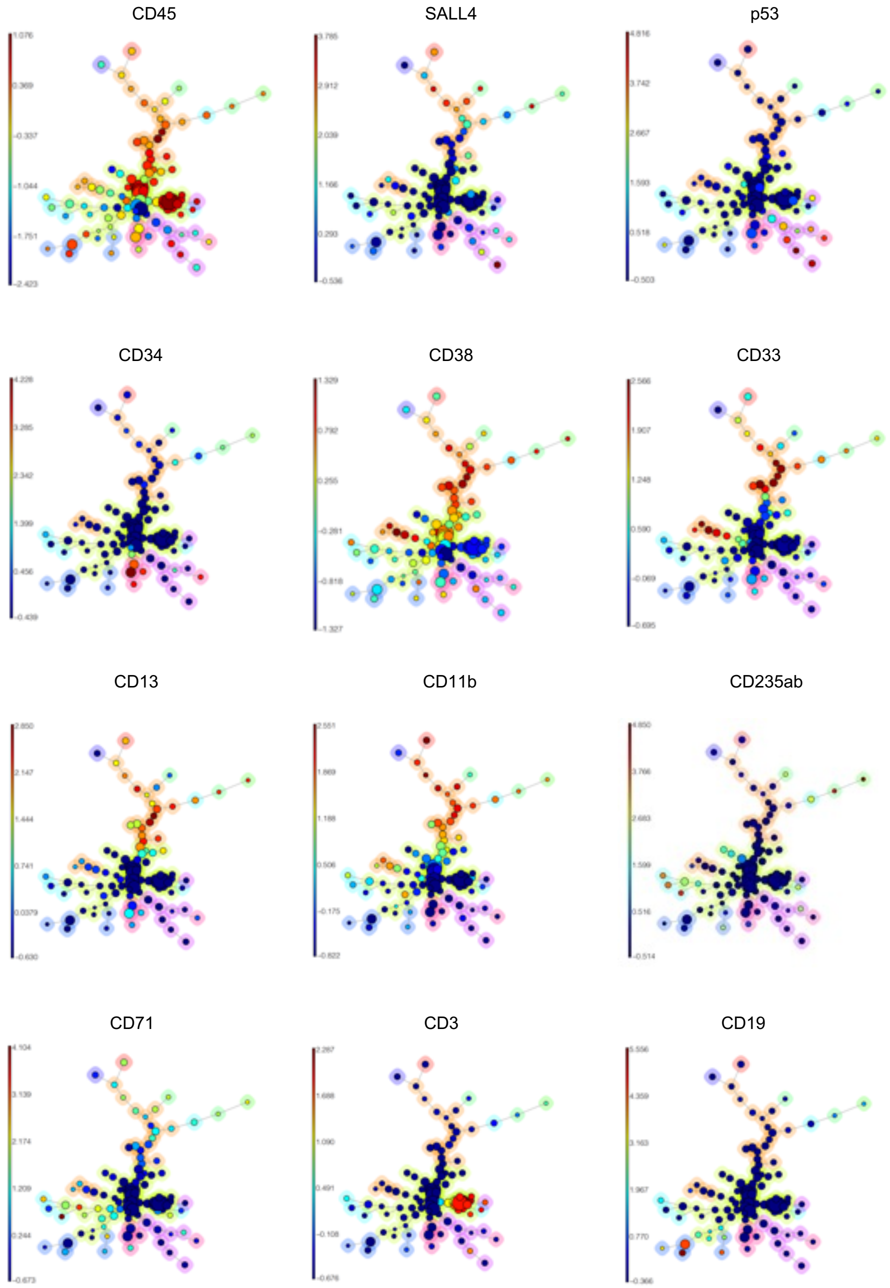


**Supplementary Figure 4. Aggregated events of surface markers**

Aggregated events of CD45, SALL4, p53, CD34, CD38, CD33 CD13, CD11b, CD235ab, CD71, CD3 and CD19.

### Supplementary Table 1. Patient clinical data

| Sample number | Age | Sex | Diagnosis | IPSS | IPSSR | Proportion of blasts in BM (%) | Hb (g/dL) | WBC ($10^3/\mu L$) | Plt ($10^3/\mu L$) |
|---|---|---|---|---|---|---|---|---|---|
| 1 | 55 | M | MDS with excess blasts2 | high | Very high | 11.6 | 7.1 | 1700 | 3.8 |
| 2 | 71 | M | Therapy-related myeloid neoplasms | Int-2 | high | 0 | 9.2 | 4200 | 2.8 |
| 3 | 65 | F | MDS with multilineage dysplasia | Int-1 | low | 0.6 | 10.2 | 2200 | 7.4 |
| 4 | 78 | F | Therapy-related myeloid neoplasms | Int-2 | high | 5.8 | 8.3 | 2300 | 2.8 |
| 5 | 63 | M | MDS unclassifiable | Int-1 | Intermediate | 0 | 9.1 | 3100 | 2.5 |
| 6 | 73 | M | MDS with excess blasts1 | Int-1 | Intermediate | 0.4 | 7.1 | 3400 | 18.8 |
| 7 | 84 | M | MDS with excess blasts1 | Int-2 | very high | 7.6% (PB blast 3%) | 8.1 | 1700 | 2 |
| 8 | 76 | F | MDS/MPN-RS-T | low | low | 0 | 8.5 | 5100 | 44.2 |
| 9 | 72 | M | MDS with multilineage dysplasia | low | low | 1.6 | 9 | 5100 | 10.9 |
| 10 | 68 | M | MDS with ring sideroblasts → AML with myelodysplasia-related changes | high | high | 21 | 6.5 | 5500 | 25.3 |

Abbreviations: MDS, myelodysplastic syndrome; WBC, Peripheral white blood cell counts; Hb, Hemoglobin; Plt, Platelets counts, IPSS, International Prognostic Scoring System; IPSSR, revised International Prognostic Scoring System; BM, bone marrow; myelodysplastic/myeloproliferative neoplasm with ring sideroblasts and thrombocytosis, MDS/MPN-RS-T.

**Supplementary Table 2. The list of Antibodies for CyTOF®**

| Antigen and Conjugate | Supplier | Catalogue Number |
|---|---|---|
| Anti-Human CD3 (UCHT1)-154Sm | Fluidigm | 3154003B |
| Anti-Human CD7 (CD7-6B7)-153Eu | Fluidigm | 3153014B |
| Anti-Human CD10 (HI10a)-156Gd | Fluidigm | 3156001B |
| Anti-Human CD11b/Mac-1 (ICRF44)-167Er | Fluidigm | 3167011B |
| Anti-Human CD13 (WM15)-147Sm | Fluidigm | 3147014B |
| Anti-Human CD19 (HIB19)-165Ho | Fluidigm | 3165025B |
| Anti-Human CD34 (581)-149Sm | Fluidigm | 3149013B |
| Anti-Human CD38 (HIT2)-144Nd | Fluidigm | 3144014B |
| Anti-Human CD41 (HIP8)-89Y | Fluidigm | 3089004B |
| Anti-Human/Mouse CD44 (IM7)-171Yb | Fluidigm | 3171003B |
| Anti-Human CD45 (HI30)-141Pr | Fluidigm | 3141009B |
| Anti-Human CD45RA (HI100)-169Tm | Fluidigm | 3169008B |
| Anti-Human/Mouse CD49F (GoH3)-164Dy | Fluidigm | 3164006B |
| Anti-Human CD56 (NCAM16.2)-163Dy | Fluidigm | 3163007B |
| Anti-Human CD90 (5E10)-159Tb | Fluidigm | 3159007B |
| Anti-Human CD117 [ckit] (104D2)-143Nd | Fluidigm | 3143001B |
| Anti-Human CD123/IL-3R (6H6)-151Eu | Fluidigm | 3151001B |
| Anti-Hu CD135/Flt3 (BV10A4H2)-158Gd | Fluidigm | 3158019B |
| Anti-Human CD235ab (HIR2)-175Lu | Fluidigm | 3175027B |
| Anti-Human HLA-DR (L243)-174Yb | Fluidigm | 3174001B |
| Anti-pAkt [S473] (D9E)-152Sm | Fluidigm | 3152005A |
| Anti-Human p53 (DO-7)-150Nd | Fluidigm | 3150024A |
| Anti-Human c-Myc (9E10)-176Yb | Fluidigm | 3176012B |
| Anti-Ki-67 (B56)-161Dy | Fluidigm | 3161007B |
| Purified anti-human CD33 | BioLegend | BL366602 |
| Purified anti-human CD71 | BioLegend | BL334102 |
| anti-SALL4 | Abnova | H00057167-M03 |
| LEAF Purified Mouse IgG1、 κ isotype Ctrl | BioLegend | BL400124 |

**Supplementary Table 3. The list of genes related myelodysplastic syndrome**

| Function | Gene |
|---|---|
| Signaling: | JAK2, CALR, MPL, CSF3R, KIT, FLT3, NF1, NRAS, KRAS, CBL, PTPN11, RIT1, GNAS, GNB1 |
| Splicing,RNA helicase: | SRSF2, SF3B1, U2AF1, U2AF2, ZRSR2, SF1, DDX41, PRPF8, LUC7L2 |
| Epigenetic: | ASXL1, TET2, DNMT3A, BCOR, EZH2, KMT2A, KMT2C, IDH1, IDH2, BCORL1, SUZ12, EED, KDM6A |
| Telomere complex: | TERT, TERC, GAR1, DKC1 |
| Transcription factor: | RUNX1, ETV6, GATA2, NPM1, CEBPA, WT1 |
| TP53, DNA repair: | TP53, PPM1D, BRCC3, CUX1, ATRX, FANCL, ATM, MYBL2 |
| Cohesins: | CTCF, RAD21, SMC3, SMC1A, STAG2 |
| Others: | SETBP1, ETNK1, PIGA |
